# Decoding emergent properties of microbial community functions through sub-community observations and interpretable machine learning

**DOI:** 10.1101/2025.08.25.671971

**Authors:** Hidehiro Ishizawa, Sunao Noguchi, Miku Kito, Yui Nomura, Kodai Kimura, Masahiro Takeo

## Abstract

The functions of microbial communities, including substrate conversion and pathogen suppression, arise not as a simple sum of individual species’ capabilities but through complex interspecies interactions. Understanding how such functions arise from individual species and their interactions remains a major challenge, limiting efforts to rationally understand microbial roles in both natural and engineered ecosystems. Because current holistic (meta-omics) and reductionist (isolation- or single-cell-based) approaches struggle to capture these emergent microbial community functions, this study explores an intermediate strategy: analyzing simple sub-community combinations to enable a bottom-up understanding of community-level functions. To examine the validity of this approach, we used a nine-member synthetic microbial community capable of degrading the environmental pollutant aniline, and systematically generated a dataset of 256 sub-community combinations and their associated functions. Analyses using random forest models revealed that the sub-community combinations of just three to four species enabled the quantitative prediction of functions in larger communities (5–9-member; Pearson’s *r* = 0.78–0.80). Prediction performance remained robust even with limited sub-community data, suggesting applicability to more diverse microbial communities where exhaustive sub-community observation is infeasible. Moreover, interpreting models trained on these simple sub-community combinations enabled the identification of key species and interspecies interactions that strongly influence the overall community function. These findings provide a methodological framework for mechanistically dissecting complex microbial community functions through sub-community-based analysis.

## INTRODUCTION

Microbial communities constitute a significant proportion of Earth’s total biomass (approximately 90 Gt-C) [1], encompassing diverse, interacting species. Far from being mere background organisms, they are key drivers of climate regulation, biogeochemical cycling, and phenotypic expression in multicellular hosts [2-4]. Their diverse metabolic capabilities are increasingly being applied in agriculture, bioproduction, and environmental remediation [5-6]. However, despite decades of research, our ability to rationally manipulate microbial community functions remains limited.

Difficulties in microbial community management largely stem from the emergent nature of microbial community functions, which are shaped by complex interspecies interactions rather than the additive effects of individual species’ capabilities [7]. For instance, in pollutant-degrading microbial communities, species lacking genetic capacity for direct degradation still influence overall system performance by supporting downstream metabolism, modulating population dynamics, or providing essential cofactors [8-10]. Existing analytical frameworks often struggle to capture such indirect contributions. Holistic meta-omics approaches provide comprehensive data but cannot easily resolve causal relationships between specific species and functional outcomes [11-12]. Reductionist approaches using isolates or single-cell data often miss the interspecies interactions that drive emergent properties. Consequently, understanding interspecies interactions and the true functional contributions of individual species remains challenging, hindering our ability to predict, design, and control microbial community functions.

A promising approach to dissect complex microbial interactions is to focus on sub-community combinations as an intermediate unit of observation. This approach involves sampling multiple combinations of a few species, typically three to five, from a larger community and evaluating their functional outputs. Unlike statistical inference from whole-community data, this approach facilitates the examination of species’ functional roles in simpler and more interpretable contexts, with still capturing emergent properties not detectable in isolates or single-cell assays. The conceptual basis for this approach derives from the theoretical food web ecology, where ecosystem-level properties have been modeled by decomposing complex systems into small ecological modules of three to four species [13-15]. Empirical studies have also shown that interactions among as few as three species can recapitulate emergent behaviors relevant to community assembly in both plant and microbial systems [16-18].

Existing studies have primarily focused on predicting species abundance or community composition, leaving open the question of how sub-community data can be leveraged to dissect functional outputs, such as substrate conversion or host phenotypic modulation. To address this gap, this study aimed to establish an analytical framework that uses sub-community information to achieve bottom-up, mechanistic understanding of microbial community functions. Specifically, a machine learning approach was applied to explain community-level functional performance based on measurements from sub-community combinations. Although previous studies have trained machine learning models on whole community data to predict functional outputs [19-20], we expected that models trained on sub-community data would further highlight influential species and interspecies interactions because of their greater simplicity and interpretability.

To evaluate the sub-community-based analytical framework in a controlled setting, we constructed a nine-member synthetic microbial community capable of degrading the environmental pollutant aniline. Aniline, a representative aromatic amine widely used to produce pesticides and dyes, is considered a high-priority bioremediation target due to its toxicity and environmental persistence [21]. Using this model system, we systematically assessed all possible sub-community combinations to test two hypotheses: (1) functional outputs of more complex communities can be predicted from simpler sub-community data, and (2) sub-community observation allows identification of key species and interspecies interactions driving emergent community functions. Although derived from a controlled synthetic system, our findings provide a methodological blueprint for leveraging sub-community data to gain mechanistic insight into complex microbial community functions.

## MATERIALS AND METHODS

### Establishment of a reproducible aniline-degrading community

To establish a model aniline-degrading microbial community, 2.5 g of soil collected from a household vegetable garden in Kakogawa City, Hyogo, Japan (34.77 N 134.85 E), was suspended in 30 mL of MSM medium [22], mixed vigorously, and centrifuged (1,000 × *g*, 1 min). Then, 5 mL of the supernatant containing microbial cells was transferred to 50 mL of MSM medium supplemented with 0.5 mM aniline (hereafter referred to as MSM aniline medium) and incubated at 30°C with shaking at 150 rpm for 48 h. Subsequently, 1 mL of this culture was inoculated into 50 mL of fresh MSM aniline medium and incubated for four days. During cultivation, 1 mL samples were periodically collected to measure residual aniline concentrations using HPLC (Table S1). The resulting enriched culture was preserved at −80°C in cryotubes containing 16.7% glycerol.

For community restoration, 1.2 mL of thawed glycerol stock was added to 10 mL of MSM medium and microbial cells were collected via centrifugation (10,000 × *g*, 4°C, 10 min), washed twice, and resuspended in 5 mL of MSM. Then, 1 mL of the cell suspension was inoculated into 50 mL of MSM aniline medium and incubated at 30°C with shaking at 150 rpm for 48 h. The resulting culture was used for subsequent analyses. To assess the community structure and aniline degradation ability, 1 mL of the culture was inoculated into 50 mL of MSM aniline medium and incubated in four replicate flasks at 30°C with shaking at 150 rpm for 48 h. Reproducibility was assessed by performing the restoration procedure twice on separate days, followed by the evaluation of residual aniline concentration using HPLC and community structure using 16S rRNA gene amplicon sequencing (methods described in Supplementary Note 1).

### Metagenome analysis

DNA was extracted from the restored aniline-degrading community using NucleoSpin Microbial DNA (Macherey-Nagel, Düren, Germany). For shotgun metagenome sequencing, libraries were prepared with the MGIEasyFS DNA Library Prep Set (MGI Tech, Shenzhen, China), and 150 bp paired-end sequencing was performed on a DNBSEQ-G400RS platform (MGI Tech) at Genome-Lead Inc. (Takamatsu, Japan). Sequencing reads were quality-filtered and adapter-trimmed with fastp v0.23.2 and assemblies were generated using metaSPAdes v3.15.5 [23]. Subsequently, reads were mapped to assembled scaffolds using BWA v0.7.17 [24], and metagenome binning was performed using MetaBATt2 v2.15 [25]. The resulting bins were evaluated using checkM v1.2.2 [26], and those with completeness >50% and contamination <5% were regarded as valid metagenome-assembled genomes (MAGs). MAGs were annotated using DFAST v1.6.0 [27] and BlastKOALA v3.0 [28]. The relative abundances of the MAGs were estimated by mapping the metagenome reads using BBMap v36.01.1. Homology search and gene feature visualizations were performed using the BLAST and CAGECAT [29].

### Isolation, identification, and characterization of synthetic community members

The restored model aniline-degrading community was diluted, spread on R2A and PCAT agar [30], and incubated at 30°C for up to one week. Morphologically distinct colonies were isolated and subjected to PCR amplification of the 16S rRNA gene using the 27F–1392R primer set [31]. Amplicons were purified using the NucleoSpin Gel and PCR Purification Kit (Macherey-Nagel) and sequenced via Sanger sequencing on a 3730xl DNA analyzer (Applied Biosystems, Foster City, CA, USA) at Macrogen Japan Corp. (Tokyo, Japan). The taxonomic positions of the isolates were identified using similarity searches against the EzBioCloud 16S database. Based on these results, nine bacterial strains were selected as members of the synthetic community: *Achromobacter veterisilvae* MA031 (Ach031), *Pseudomonas monteilii* MA033 (Pse033), *Sphingomonas pseudosanguinis* MA034 (Sph034), *Microbacterium laevaniformans* MA039 (Mic039), *Bosea spartocytisi* (Bos046), *Bacillus mexicanus* MA050 (Bac050), *Pseudomonas* sp. MA056 (Pse056), *Caballeronia cordobensis* MA074 (Cab074), and *Sphingomonas* sp. MA075 (Sph075) (Table S2). Given the absence of rRNA gene sequences in our MAG sequences, we predicted the closest MAG–isolate correspondence by comparing species-level classifications obtained independently for MAGs and isolates. The nine strains were cultivated overnight in 2 mL R2A medium in a 24-well plate (30°C, 150 rpm), washed twice with MSM medium, and used for subsequent experiments.

For one isolate (Cab074), identified as the primary aniline degrader, a fluorescence marker (eGFP) was introduced using the pMRE152 plasmid as previously described [32]. The obtained fluorescent transformants (hereafter Cab074F) were purified and used as the member of synthetic communities in place of the wildtype.

To evaluate the ability of each synthetic community member to utilize aniline, individual strains were inoculated into 500 µL MSM aniline medium in a 24-well plate at an initial OD_600_ of 0.01. After 48 h of incubation, the residual aniline content was quantified using HPLC.

### Full factorial evaluation of the sub-community combinations

A 384-well plate-based assay was conducted to evaluate all possible sub-community combinations of the synthetic community that included Cab074F (i.e., 2□ = 256 patterns; *n* = 4). Each well of the black 384-well plate contained 20□µL of MSM aniline medium supplemented with 2□µg/mL of resazurin, a metabolic marker reduced to fluorescent resorufin in proportion to microbial aerobic metabolism [33]. Because aniline was the sole carbon and energy source, fluorescence intensity was considered a proxy for aniline utilization. Strains were inoculated into wells at an initial OD□□□ of 0.001 each. To minimize pipetting errors, a robotic pipettor (Assist Plus, Integra Biosciences, Zizers, Switzerland) was used for culture preparation. The plates were incubated in a microplate reader (Synergy HTX, Agilent Technologies, Santa Clara, CA, USA) at 30°C with orbital shaking at 237 rpm. During the 72 h cultivation period, GFP (excitation/emission: 485/528 nm) and resorufin (528/590 nm) fluorescence were measured every 10 min. Three plates were analyzed to cover all sub-community combinations in four replicates each. Every plate included at least 16 Cab074F-only and 16 blank wells without microbial inoculation.

The “aniline-utilization activity” and the “Cab074F growth” in each well were defined as the area under the curve (AUC) calculated from the fluorescence time courses of resorufin and GFP during cultivations, respectively. The detailed calculation method is described in Supplementary Note 2. We also checked the correlation between aniline degradation rate and resorufin fluorescence, as well as between colony-forming unit of Cab074F and GFP fluorescence, using selected combinations (Supplementary Note 3).

The functional effect of each non-degrader strain was calculated as the change in aniline-utilization activity between sub-communities that differed only by the presence or absence of the focal strain. The Shapley value, representing the overall contribution of each species to community function, was calculated as the average functional effect across all possible assembly pathways [34]. The functional interactions between all pairs of non-degrader strains were estimated as the difference in the mean functional effect of a focal strain in the presence versus absence of a partner strain. Statistical significance of differences in functional effect was assessed using two-sided *t*-test in R v4.4.2.

### Random forest model

Random forest models were employed to examine the bottom-up predictability of aniline-utilization activity. Models were constructed using the “randomForest” R package, with the presence–absence data of eight non-degrader strains as explanatory variables and the measured aniline-utilization activity as the response variable. We chose species presence/absence rather than their abundances, because species abundances in microbial communities are highly dynamic, and relying on single snapshot can introduce sampling bias that does not reflect overall system performance.

To determine which levels of sub-community diversity (species richness) carry important information for predicting community-level functions, we first trained models with observed sub-community combinations at fixed richness levels: two (n = 8), three (n = 28), four (n = 56), five (n = 70), six (n = 56), and seven-member combinations (n = 28). Prediction performance was then evaluated as Pearson’s correlation coefficient using data from 5–9-member combinations (n = 163) as the test set. To prevent data leakage, when a test combination is also present in the training pool, we rebuilt the model after removing only that single overlapping data point from the training data. Furthermore, to assess model performance under limited training data, models were also trained on using randomly sampled subsets comprising 50% or 25% of the original training set.

To identify important species and interspecies interactions contributing to prediction accuracy, permutation importance and Friedman’s H statistic were extracted from the random forest models using the “vivid” R package [35]. Models trained on all sub-community combinations at each richness level (two-to six-members) and a model trained on all 2–4-member sub-communities were used for this analysis. The obtained values were compared with empirically calculated Shapley value and functional interaction strength to evaluate their ability to identify key species and interactions.

To account for variability in training and sampling, the entire procedure (training and evaluation) was repeated 100 times with different random seeds; we report the mean and standard deviation across runs. Figures and parameter inference were generated from the model with median performance (51st of 100 trials).

### Data availability

The 16S rRNA gene sequences of the bacterial isolates were deposited in the DDBJ/EMBL/GenBank database under accession numbers LC878797-LC878805. Metagenome sequence reads were deposited under the accession number DRR707044. All experimental data and analytical codes have been deposited in Zenodo [36].

## RESULTS

### Reproducibility of the model aniline-degrading microbial community

A major challenge in systematic analysis of microbial communities is the lack of reproducibility (i.e., the ability to regenerate the same community across experiments). To address this, we first established a reproducible community using a whole-community glycerol stock derived from an aniline-degrading community. Restoring the preserved stock through a standardized procedure confirmed consistent community degradation of 0.5□mM aniline within 48 h and a stable community structure dominated by specific genera including *Achromobacter, Burkholderia*, and *Pseudomonas*, with only minor variations in relative abundances (Fig. S1). The dominant bacteria in the restored communities were consistent at the ASV level, suggesting that the preserved community stock provides a reliable source of reproducible model aniline-degrading communities.

### Sequence-based prediction of aniline- and catechol-degrading bacteria

Shotgun metagenome analysis was performed to identify species in the aniline-degrading community harboring genes responsible for aniline degradation. The assembly of approximately 107 million reads reconstructed 13 MAGs, encompassing all dominant taxa detected via the 16S rDNA amplicon sequencing (Fig. 1A; Table S3).

**Fig. 1.**
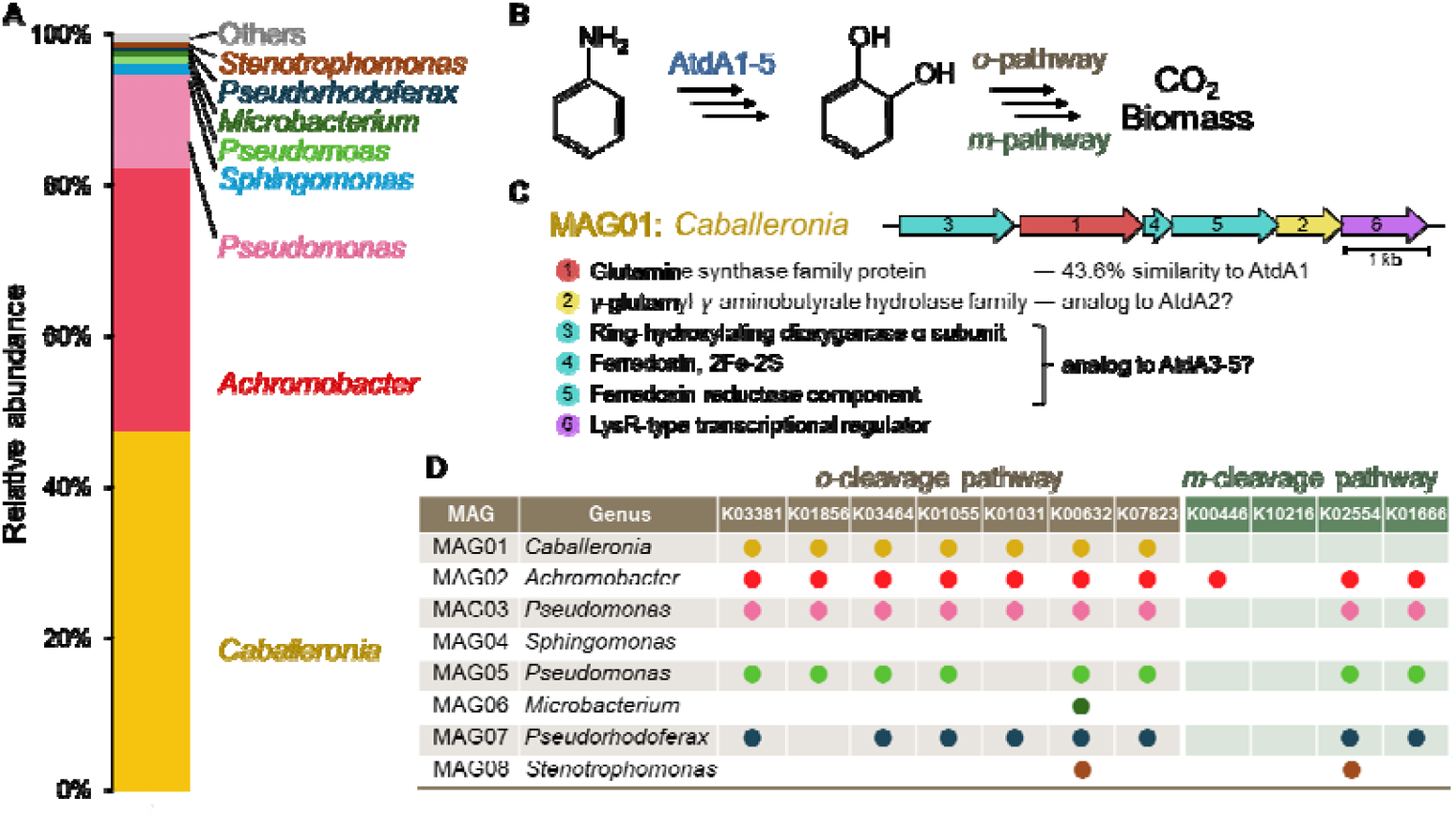
Metagenomic characterization of the model aniline-degrading community served as the model system in this study. A) Community composition of the aniline-degrading community. Relative abundances of metagenome-assembled genomes (MAGs) and their closest genera are shown. B) Schematic representation of the known aniline degradation pathway. AtdA1-5 catalyze the initial oxidation of aniline, and the intermediate catechol is degraded via the ortho- or meta-cleavage pathway. C) The putative aniline oxidation gene cluster identified in MAG01. Annotation of the six gene products and their sequence or functional similarity to known aniline oxidation enzymes are shown. D) Distribution of catechol-degradation genes among the eight most abundant MAGs in the community. Presence/absence of KEGG entries associated with the catechol ortho- and meta-cleavage pathway is indicated.

Reportedly, aniline degradation is initiated by a unique reaction in which a glutamate synthase-like protein (AtdA1) attaches a glutamate molecule to aniline, forming γ-glutamylanilide. Subsequently, dioxygenase proteins (AtdA3–5) oxidize the aromatic ring, yielding catechol, glutamate, and ammonium [37-38] (Fig. 1B). Genes encoding these enzymes are often clustered with *atdA2*, encoding the reverse reaction of AtdA1. Catechol, a common intermediate in aromatic compound biodegradation is typically introduced into central anabolic metabolism via either the ortho- or meta-cleavage pathways [39].

Within the obtained MAGs, only MAG01, assigned to the genus *Caballeronia*, contained a gene with significant similarity to known *atdA1* genes (43.6% amino acid sequence similarity to AtdA1 of *Acinetobacter* sp. YAA) [37]. Adjacent genes encoded components of an aromatic ring-hydroxylating dioxygenase (a functional analog of AtdA3–5) and a glutamyl hydrolase (a functional analog of AtdA2) (Fig. 1C). Although these surrounding genes exhibited <20% similarity to previously reported aniline-degrading gene clusters, we considered this gene cluster responsible for aniline degradation in our community.

For catechol degradation, several MAGs dominant in the community—including those affiliated with *Caballeronia, Achromobacter, Pseudomonas*, and *Pseudorhodopherax*—possessed most genes involved in the ortho-cleavage pathway (Fig. 1D). In contrast, no MAG carried the candidate genes for 2-keto-4-pentenoate hydratase (K02554), a key enzyme in the meta-cleavage pathway, indicating that catechol degradation in this community proceeds primarily through the ortho-cleavage pathway.

In summary, the metagenomic analysis suggests a potential metabolic relay within the model aniline-degrading community, in which *Caballeronia* first converts aniline into catechol, whereas several other strains contribute to its subsequent metabolism. Bacteria lacking the ability to degrade neither aniline nor catechol may persist through metabolic cross-feeding or scavenging of cell debris.

### Characterization of synthetic community members

To construct a tractable synthetic community reflecting the diversity, functionality, and ecological characteristics of the model aniline-degrading community, nine bacterial strains representing a broad range of phylogenetic groups present in the original community were isolated and selected as members of the synthetic community (Table S2). Species-level taxonomic comparisons indicated that Cab074, Ach031, Pse033, Sph075, Pse056, and Mic039 corresponded most closely to the six dominant MAGs (MAG01–MAG06). In contrast, Sph034, Bos046, and Bac050 were considered minor community members without corresponding MAGs. To facilitate detection of the primary aniline degrader, a GFP-labeled strain of Cab074 (Cab074F) was constructed and used in place of the wildtype in subsequent experiments. Degradation assays for individual synthetic community members confirmed that only Cab074F degraded aniline (Fig. S2), consistent with the metagenome characterization of the original community. Overall, the synthetic community recapitulates key characteristics of the original community: a single *Caballeronia* genotype predominantly performs initial aniline oxidation and other strains would participate in downstream metabolism.

### Composition–function landscape

Using the synthetic community, we investigated how community-level functions emerge from interactions between the aniline degrader and non-degrader strains, using a microplate-based assay (Fig. 2A). To increase throughput, we employed a fluorescence-based assay of aniline-utilization activity using resazurin. Preliminary tests showed a positive correlation between RFU and aniline degradation, as evaluated using HPLC (Pearson’s *r* = 0.77) (Fig. S3). Although the correlation was moderate, deviations likely arose because resazurin reduction reflects not only aniline degradation but also the metabolism of downstream intermediates. Thus, the resazurin-based assay would capture overall aniline-utilization activity, encompassing both aniline degradation and downstream metabolic processes. Simultaneously, Cab074F abundance was monitored by using GFP fluorescence, which showed strong linear correlation between GFP RFU and MA074F abundance as determined via colony counting (Pearson’s *r* = 0.89) (Fig. S3).

**Fig. 2.**
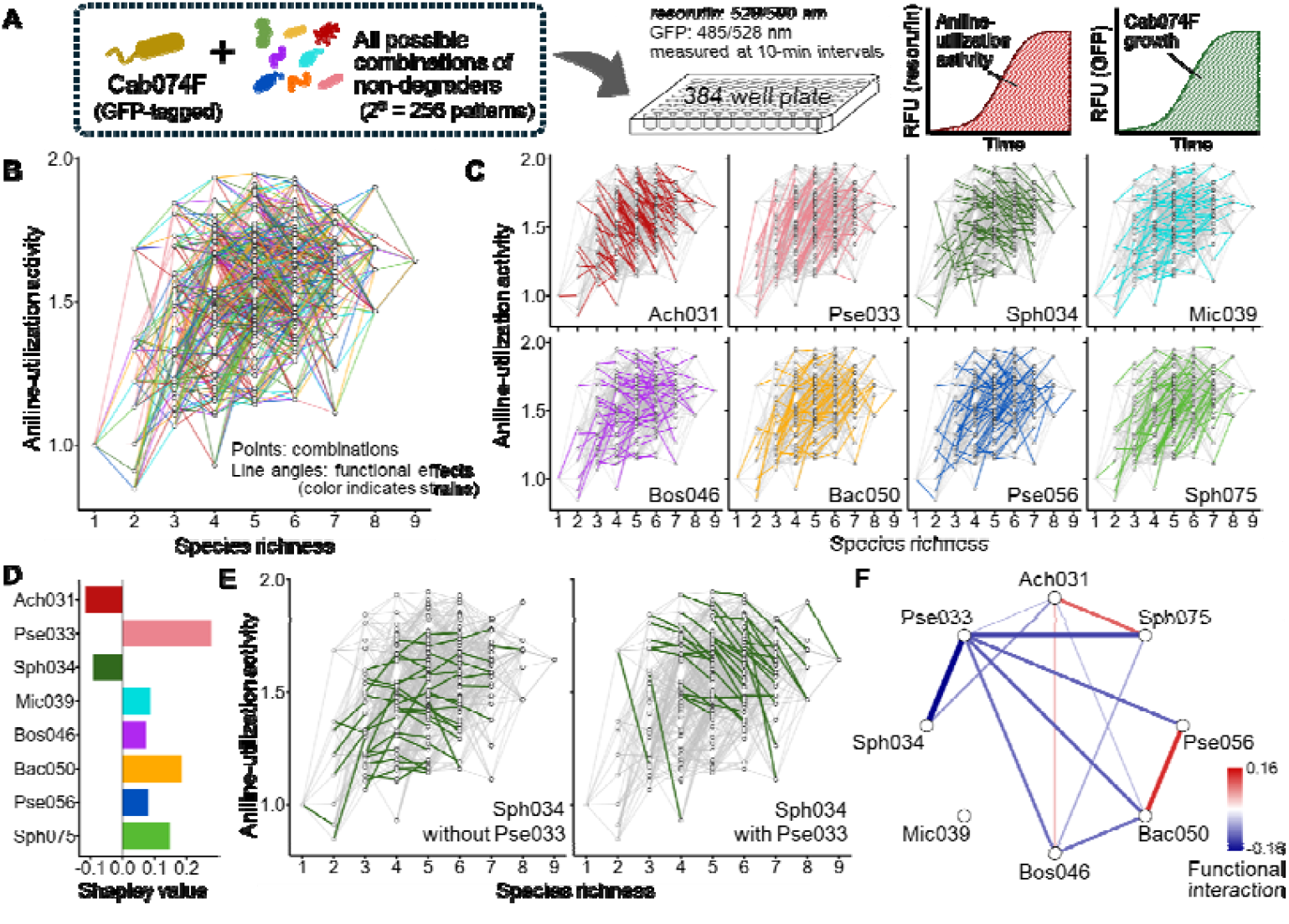
Full factorial evaluation of the synthetic aniline-degrading community. A) Schematic overview of the experimental workflow. Sub-community combinations consisting of the primary aniline degrader (Cab074F) and all possible combinations of eight non-degrader strains were cultivated for 72 h in medium containing aniline as the sole carbon source. Aniline-utilization activity and Cab074F growth were quantified as the area under the fluorescence-time curve of resorufin and GFP, respectively. B) Composition–function landscape of aniline-utilization activity for all 256 combinations. Each data point represents aniline-utilization activity of a sub-community combination at a given species richness. Line angles reflect the functional effect of each strain on aniline-utilization activity, and colors indicate the identity of the strain present in the right-side node but absent in the left-hand node (color-coded as in panel C). C) Strain-resolved composition–function landscapes for eight non-degrader strains. D) Overall contributions of the eight non-degrader strains to community-level function, quantified as Shapley values. E) Example of a functional interaction: Sph034 exhibited a near-neutral effect in the absence of Pse033 but a negative effect when Pse033 was present. The change in the mean functional effect was defined as the functional interaction between two species. F) Pairwise functional interactions among the eight non-degrader strains. Positive values indicate the synergistic contributions to community-level functions; negative values indicate sub-additive contributions. Statistically significant interactions (t-test, *P* < 0.05) are indicated.

By quantifying the aniline-utilization activity of all possible sub-community combinations containing Cab074F (i.e., 2□ = 256 patterns), we generated a composition–function landscape in which each data point represents a sub-community combination and each line denotes the functional effect of a single non-degrader (Fig. 2B). In this study, “functional effect” was defined as the change in aniline-utilization activity between two sub-community combinations differing only in the presence or absence of the focal strain. If the functional effect of each strain were constant regardless of co-existing members, the line angles would remain consistent. Indeed, Ach031 and Pse033 consistently exhibited negative and positive functional effects, respectively, with few exceptions (Fig. 2C). Sph034 and Mic039 also showed relatively consistent effects, whereas the remaining strains exhibited highly variable effects, ranging from strongly positive to strongly negative depending on the composition of the surrounding community.

To systematically quantify the contribution of each species to community-level function, we applied Shapley values to the composition–function landscape, calculating the average functional effect of each strain across all possible assembly pathways from the Cab074F-only community to the nine-member community (8! = 40,320 pathways). As the results, Pse033 contributed most positively, followed by Bac050 and Sph075, whereas Ach031 and Sph034 contributed negatively (Fig. 2D).

We evaluated how the presence or absence of a given non-degrader strain altered the functional effect of another. For example, Sph034 had a near-neutral effect in the absence of Pse033, but became negative when Pse033 was present (Fig. 2E). We quantified such “functional interaction” as changes in the mean functional effect of a focal strain due to a partner strain’s presence or absence. We identified ten significant negative and three positive functional interactions (*t*-test, *P* < 0.05), indicating combinations that lead to either diminished or enhanced community-level functions, respectively, compared with simple additive assumptions (Fig. 2F). Despite its high individual contribution to aniline degradation, Pse033 exhibited negative interactions with six other strains, suggesting diminished contributions in co-culture. In contrast, Mic039 showed no significant interactions, indicating that its functional effect was largely independent of other species.

### Abundance of primary aniline degrader

Given that Cab074F was the sole strain capable of converting aniline to catechol, we hypothesized that variations in its population success could explain differences in community-level aniline-utilization activity. To test this hypothesis, we generated a composition–abundance landscape for Cab074F using GFP fluorescence data (Fig. S4) and assessed the correlations between the functional effect on aniline-utilization activity and the corresponding effects on Cab074F growth within the same community contexts (Fig. 3). The results varied significantly across non-degrader strains: Ach031, Pse033, Sph034, and Mic039 showed significant positive correlations, whereas the others showed weak or no correlations. These positive correlations suggest that the influence of these non-degrader strains on community-level aniline-utilization activity can be largely explained by their facilitation or inhibition of Cab074F growth. In contrast, the absence of correlation implies the involvement of additional mechanisms, such as effects on downstream metabolism or interactions with coexisting non-degraders. It was found that Ach031 and Pse033, which exhibited consistent functional effects regardless of community composition (Fig. 2C), also showed exceptionally strong positive correlations. In contrast, strains that exhibited highly variable functional effects (Bos046, Bac050, Pse056, and Sph075) showed weak or no correlations. These findings suggest that the functional effects mediated through Cab074F growth tend to be robust to community composition, whereas effects involving other mechanisms are more sensitive to the community context.

**Fig. 3.**
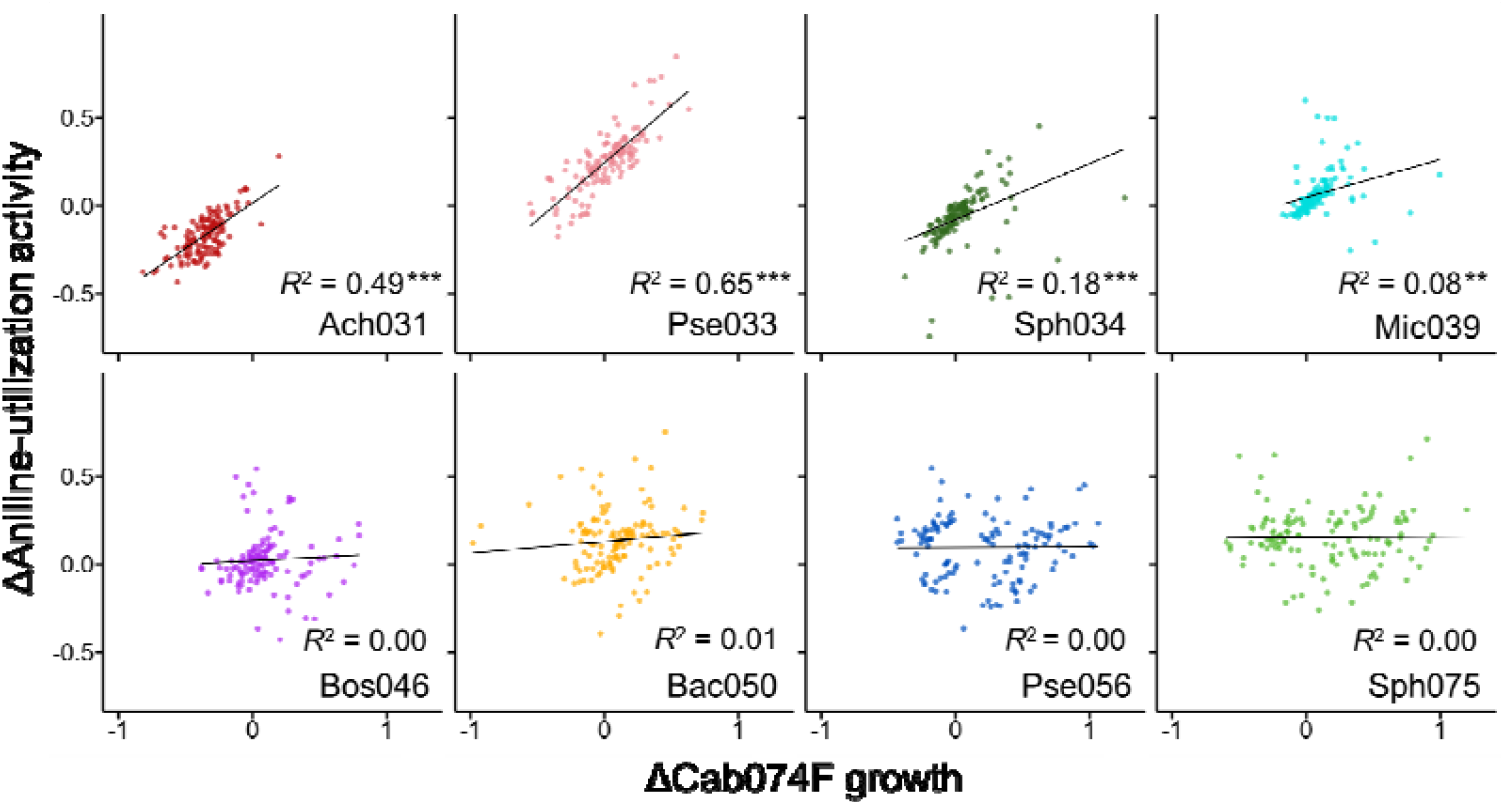
Correlation between changes in Cab074F’s growth and aniline-utilization activity. Each point represents differences in Cab074F growth (measured using GFP) and aniline-utilization activity (measured using resazurin) between sub-community combinations differing only by the presence or absence of a focal non-degrader strain (corresponding to a line in Fig. 2C). A positive correlation suggests that each species’ functional effect is related to promoting or inhibiting the growth of the primary aniline degrader, Cab074F, whereas the absence of a correlation implies that other mechanisms predominantly contribute. Asterisks denote significance levels in Pearson’s correlation analysis: ^***^ *P* < 0.001; ^**^ *P* < 0.01; ^*^ *P* < 0.05.

### Bottom-up predictability of the composition-abundance landscape

Using the obtained dataset, we tested whether the functions of complex microbial communities can be predicted based on observations of simpler sub-community combinations. To assess how species richness in the training data influences predictive performance, we trained random forest models using sub-communities of defined richness levels and applied them to predict aniline utilization activity in communities containing 5–9 species. Models trained solely on three-member combinations (n = 28) achieved high predictive accuracy for more complex communities (*r* = 0.78) (Fig. 4A). Incorporating more diverse combinations into the training set did not substantially improve the prediction performance (Fig. 4B). However, models trained solely on two-member combinations (i.e., Cab074F plus one other strain, n = 8) showed markedly lower accuracy (*r* = 0.49). These results indicate that sub-community combinations of just three to four species can support robust bottom-up prediction of microbial community functions.

**Fig. 4.**
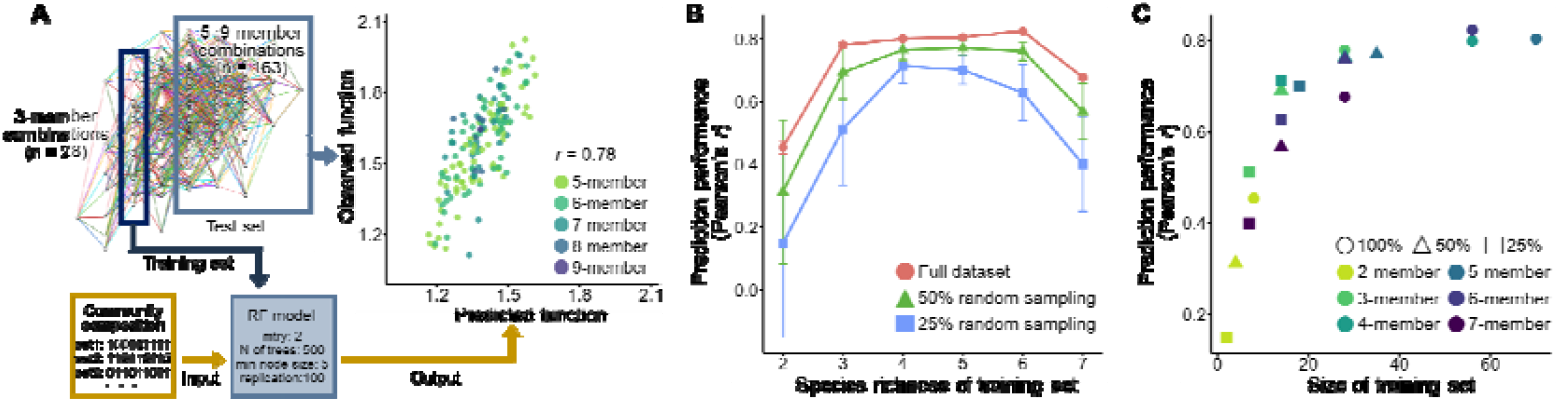
Bottom-up prediction of aniline-utilization activity using random forest models. A) Schematic of the bottom-up prediction framework. To understand which levels of sub-community diversity (species richness) are informative for predicting community-level functions, random forest models were trained on observed sub-community combinations at fixed richness levels (example shown for species richness = 3). Model performance in predicting aniline-utilization activity of 5–9-member combinations was evaluated as Pearson’s correlation coefficient. B) Prediction performance of random forest models trained on sub-community combinations of varying species richness. To assess model performance under limited training data, models trained on reduced datasets (50% or 25% random sampling) were also evaluated. Error bars indicate the standard deviations across 100 runs. C) Relationship between prediction performance and the number of sub-community combinations used for model training. Each data point represents a model trained at the indicated species richness and saturation level.

Because exhaustively sampling all sub-community combinations is infeasible in real microbial communities, we tested performance with reduced training data (50% and 25%). Reducing the dataset caused only modest decreases in predictive performance. For example, models trained on 25% of the three-member combinations (n = 7) exhibited lower performance (*r* = 0.51), whereas those trained on 25% of the four-member combinations (n = 14) still achieved high performance (*r* = 0.71) (Fig. 4B). Across conditions, prediction performance was tightly associated with the number of unique sub-community combinations used for training (Fig. 4C), explaining the apparent dip in performance at species richness of 7, where relatively few combinations are possible (n = 28). Moreover, when comparing models at similar training set sizes, those trained on 3–4-member combinations tended to outperform models trained on more diverse (6–7-member) combinations. Overall, random forest models trained with simple sub-community combinations can reliably predict functions of diverse communities without exhaustive data, provided that the training set included the sufficient number of unique combinations.

### Interpretation of the random forest model to identify key species and interactions

We tested whether interpretating the trained random forest models could reveal key species and interspecies interactions that shape community-level functions. To this end, we evaluated two model-derived metrics: permutation importance, which quantifies each feature’s contribution by measuring changes in prediction accuracy after random shuffling of focal features [40] and Friedman’s H statistic, which estimates the strength of interactions between features by assessing how the effect of one feature on aniline utilization activity depends on another [41].

We applied these interpretability techniques to the random forest model trained on three-, four-, and five-member sub-community combinations (Fig. 5A–C). The permutation importance scores consistently identified Ach031 and Pse033 as the most influential species, followed by Sph034, Bac050, and Sph075. This ranking closely matched species contributions estimated from the full dataset using Shapley values (Fig. 2D), indicating that random forest models trained on simple sub-community data can reliably identify the dominant contributors to community-level functions.

**Fig. 5.**
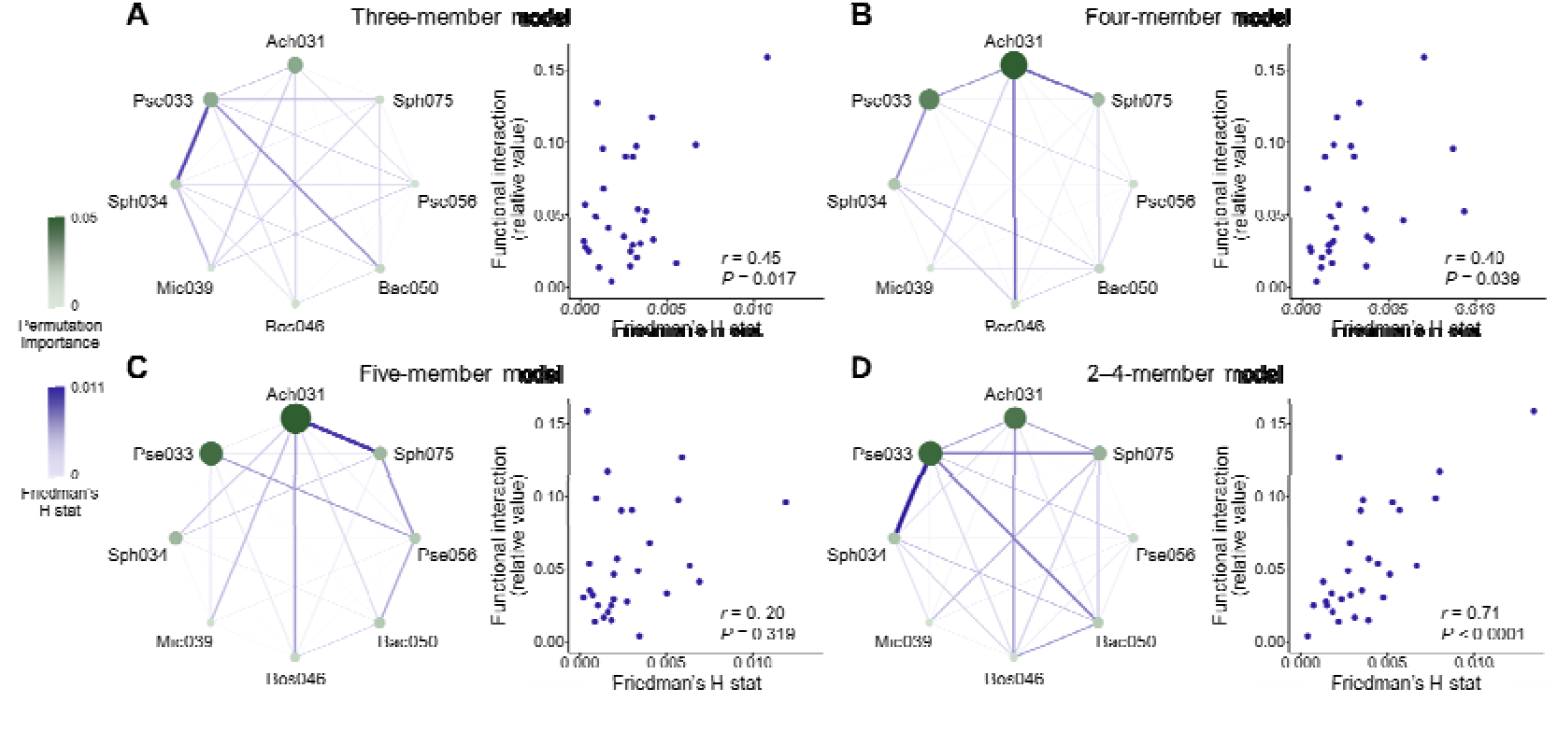
Key species and interspecies interactions inferred from the random forest models. Permutation importance and Friedman’s H statistic were calculated from random forest models trained on sub-community combinations comprising three, four, and five members, as well as from models trained on two- to four-member combinations. Permutation importance was used to quantify the relative contributions of individual species to community-level aniline utilization activity, whereas Friedman’s H statistic was used to assess the strength of pairwise functional interactions. To evaluate the reliability of inferred interactions, Friedman’s H values were compared with the empirically derived functional interaction strengths (Fig. 2F). Permutation importance values were consistent with empirically calculated Shapley values (Fig. 2D).

Additionally, Friedman’s H statistics derived from models trained on three- and four-member sub-communities correlated significantly with empirical functional interaction strengths (Fig. 2E) (*r* = 0.40–0.45, *P* < 0.05), whereas those trained on two-member sub-communities showed no correlation (*r* = 0.14, *P* = 0.457). The three- and four-member models highlighted different sets of pairwise relationships, and predictive accuracy improved markedly when models were trained on combined dataset of all two- to four-member combinations (*r* = 0.71, *P* < 0.0001) (Fig. 5D). In contrast, Friedman’s H statistics derived from models trained on five-member sub-communities were no longer significantly correlated with the observed functional interactions (*r* = 0.20, *P* > 0.05), and a similar loss of correspondence occurred in six-member models (*r* = 0.14, *P* > 0.05). These results suggest that increasing the species richness of training data reduces the ability to resolve specific interspecies interactions, likely due to statistical difficulties in pinpointing influential feature pairs within high-dimensional observational units.

Together, these findings show that machine-learning models trained on simple sub-community combinations can identify key species and interspecies interactions underlying emergent community functions. In particular, sub-community combinations of three to four species appear to represent the most informative unit for dissecting the emergent properties of microbial community functions.

## DISCUSSIONS

Using a synthetic microbial community capable of degrading aniline, this study showed that analyzing simple sub-community combinations enables the identification of key species and interactions shaping community-level functions (Fig. 5). Traditional analytical approaches to microbial community analysis have tended to follow either holistic (meta-omics-based) or reductionist (isolated strain/single-cell-based) strategies. In contrast, this study proposes an intermediate, sub-community-based framework that effectively captures the emergent properties of microbial community functions. Our proposed framework involves three steps: (1) generating multiple sub-community combinations from a target microbial community, (2) evaluating their functions and compositions, and (3) analyzing the resulting dataset to infer the species contributions and interactions, thereby identifying rational targets for follow-up analyses or microbial community engineering. Combining with meta-omics approaches (e.g., metatranscriptomics), this framework would further strengthen mechanistic inference. Recent advances in high-throughput microbial cultivation platforms [42-43] have made it increasingly feasible to generate and characterize thousands of sub-community combinations without isolation. Thus, the findings and analytical framework presented here provide a foundation for applying this approach to more complex natural microbial communities in the near future.

Consistent with recent studies investigating composition–function landscapes of synthetic microbial communities [44-45], our results demonstrate that the functional effects of individual species are highly context-dependent (Fig. 2). To explain such context dependency, a “global epistasis” has been proposed in which a species’ functional effect can be approximated by simple linear regression against the functional level of the background community—tending to be positive when the baseline function is low, and negative when it is high [7]. This pattern was evident in our dataset (Fig. S5) and appears to reflect both functional redundancy among species and complex interaction networks that obscure individual contributions. Beyond this statistical explanation, our results suggest that mechanistic differences in species’ modes of action also contribute: strains that affected aniline degradation through direct promotion of the degrader Cab074F’s growth showed relatively stable functional effects, whereas those acting through other mechanisms exhibited greater context sensitivity (Fig. 3). Although detailed interaction mechanisms are beyond the scope of this study, linking statistical patterns in composition–function landscapes with mechanistic insights constitutes a future avenue for effectively controlling microbial community functions.

Another key finding is that microbial community functions can be predicted in a bottom-up manner by combining simple sub-community observations with machine learning. Similar bottom-up approaches have recently gained traction in studies using synthetic microbial communities, with several strategies showing promise. For example, genome-scale metabolic models have been applied to predict the metabolic output of complex microbial consortia [46-48]. In parallel, mathematical models that infer community function from the geometry of composition–function landscapes have also been developed [7,45,49], enabling prediction without detailed knowledge of metabolic pathways. Although many of these modeling studies relied on full datasets in composition–function landscapes and leave-one-out validation strategies, our framework demonstrated that reasonable prediction can be achieved using only a limited set of simple sub-community combinations. This flexibility makes our approach suitable not only for the rational design of synthetic microbial consortia but also for investigating natural communities, where exhaustive sub-community observation is infeasible.

This study also introduced a method for identifying key species and interspecies interactions by interpreting variable-importance measures derived from machine-learning models. Whereas models trained on 3–4-member communities yielded reasonable predictions, increasing species richness in observed sub-community combinations did not necessarily improve analytical insights (Fig. 5), likely due to statistical challenges in resolving interspecies interactions from more complex combinations. These findings suggest that sub-community combinations consisting of three to four species may represent an analytical sweet spot, balancing predictive power and interpretability. This observation aligns with previous reports where three- to four-member sub-communities provided robust models for predicting species abundance or survival [17-18], highlighting their promise as a unit for dissecting emergent properties in microbial communities.

In summary, this study demonstrates that simple sub-community combinations contain valuable information for bottom-up, mechanistic understanding of microbial community functions. The machine learning–based framework proved robust to data sparsity, making it a promising tool for analyzing diverse and noisy microbial systems. Although further validation is needed to assess its applicability to highly diverse and dynamic microbial ecosystems, integrating this sub-community-based strategy with existing meta-omics and cultivation-based methods offer a promising path towards rationally predicting and controlling emergent properties of microbial community functions.

## ACKNOWLEDGEMENTS

This work was supported by JST, ACT-X (JPMJAX22B1), Research Grants by Kurita Water and Environment Foundation (22E053, 23K008), and Gene-grant 2023 by Nippon Genetics Co., Ltd. to Hidehiro Ishizawa. The manuscript’s English was polished using generative AI for grammar and style and further edited by Enago (www.enago.jp). The authors reviewed and edited all output.

## AUTHOR CONTRIBUTIONS

Hidehiro Ishizawa: Conceptualization, Formal analysis, Funding acquisition, Project administration, Software, Visualization, Writing – Original Draft. Sunao Noguchi: Formal analysis, Investigation, Visualization, Writing – Review & Editing. Miku Kito: Investigation, Writing – Review & Editing. Yui Nomura: Investigation, Writing – Review & Editing. Kodai Kimura: Investigation, Writing – Review & Editing. Masahiro Takeo: Project Administration, Writing – Review & Editing, Supervision.

