## Supplementary Files for "Decoding emergent properties of microbial community functions through sub-community observations and interpretable machine learning"

**Supplementary Note 1**

For amplicon sequencing, the 16S rRNA gene library (515F–806R) was prepared using a two-step PCR procedure and sequenced on the Illumina Miseq (2×300 bp) at Bioengineering Lab. Co. Ltd. (Sagamihara, Japan). TaKaRa Ex Premier DNA Polymerase (Takara Bio, Shiga, Japan) was used for PCR according to the manufacture’s protocol. Sequencing reads were processed using DADA2 [1] to generate an ASV table, and taxonomy was assigned using the SILVA v138 database [2].

**Supplementary Note 2**

AUC was calculated using the R package “gcplyr” [3] as follows: (1) baseline correction by subtracting the mean relative fluorescence units (RFU) values of blank wells on the same plate; (2) alignment of time windows across plates by shifting to match the lag phase of the Cab074F-only wells, as estimated using gcplyr; (3) normalization of the RFU values by setting adjusted time 0 to 1 for each well; and (4) normalization of the AUC values by dividing the mean AUC of the Cab074F-only wells.

**Supplementary Note 3**

To validate that resorufin fluorescence reflected aniline-utilization capacity, cultivation assays for 18 randomly selected combinations were performed using the same procedure to the 384-well plate-based assay but scaled up to 300 µL in a 96-well plate to obtain sufficient samples for HPLC. Cultures were prepared in duplicate wells, and samples were collected after incubating 48 and 72 h to evaluate aniline degradation via HPLC. These results were compared with the RFU values of resorufin measured at the corresponding time points. Similarly, the correlation between GFP fluorescence and Cab074F abundance was validated by plating 13 Cab074F cultures in a 96-well plate after 24, 48, and 72 h on MSM aniline medium solidified with agarose and comparing Cab074F abundance in CFU/mL with the corresponding GFP RFU values.

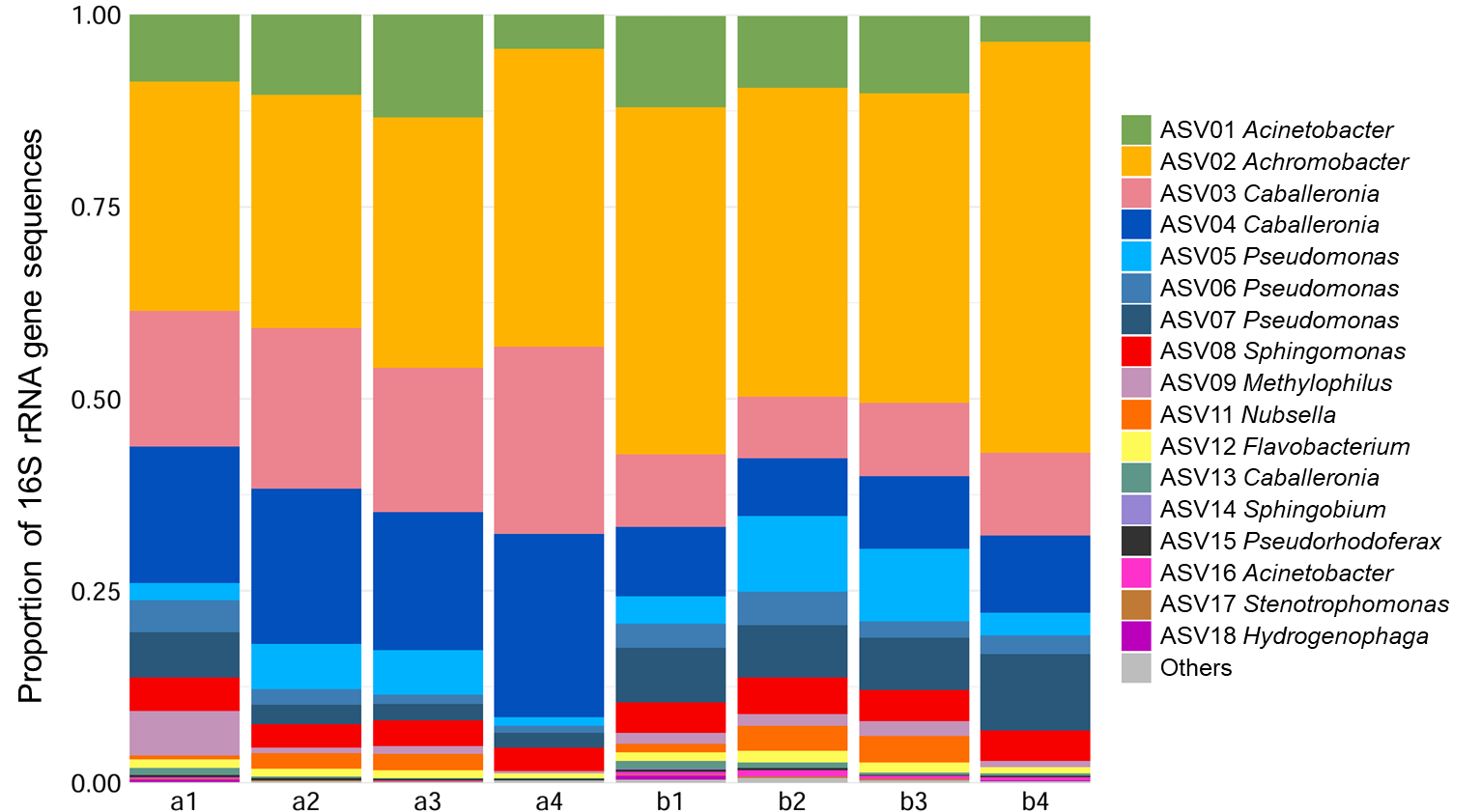

**Fig. S1** Reproducibility of community composition in the restored model aniline-degrading community. The community was restored twice from glycerol stocks on different dates (labels “a” and “b” denote the two restoration dates), each in four replicates (see Materials and Methods). Community composition at 48 h was assessed by amplicon sequencing.

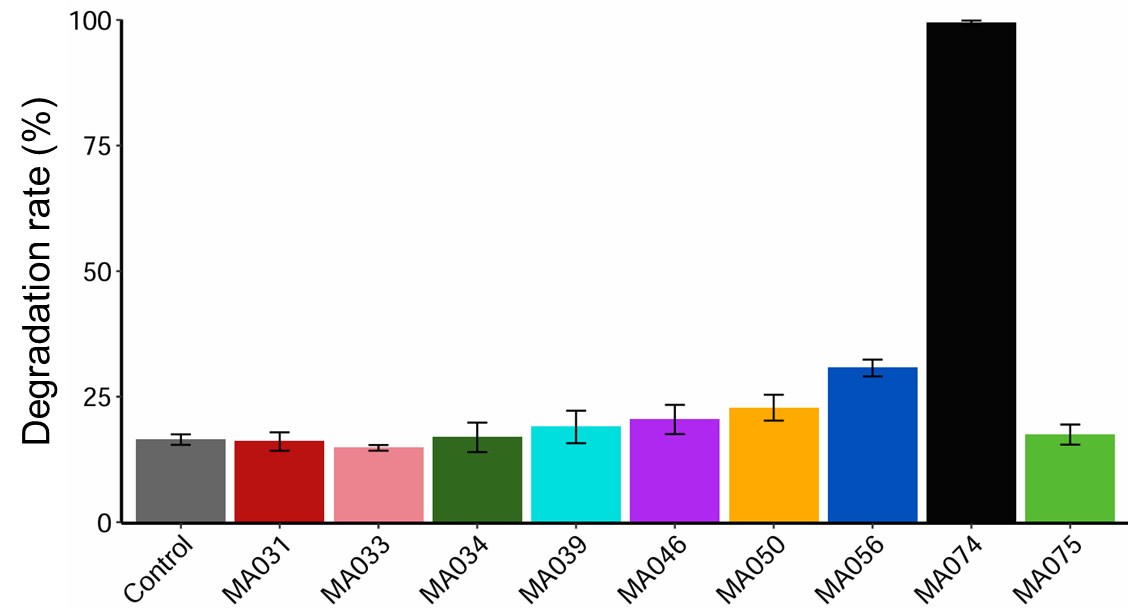

**Fig. S2** Aniline-degrading ability of the nine synthetic community members. Degradation rates for 2 mM aniline over 48 h of cultivation are shown. The observed decrease in the　uninoculated control reflects evaporation during cultivation. Error bars represent the standard deviation (n = 3).

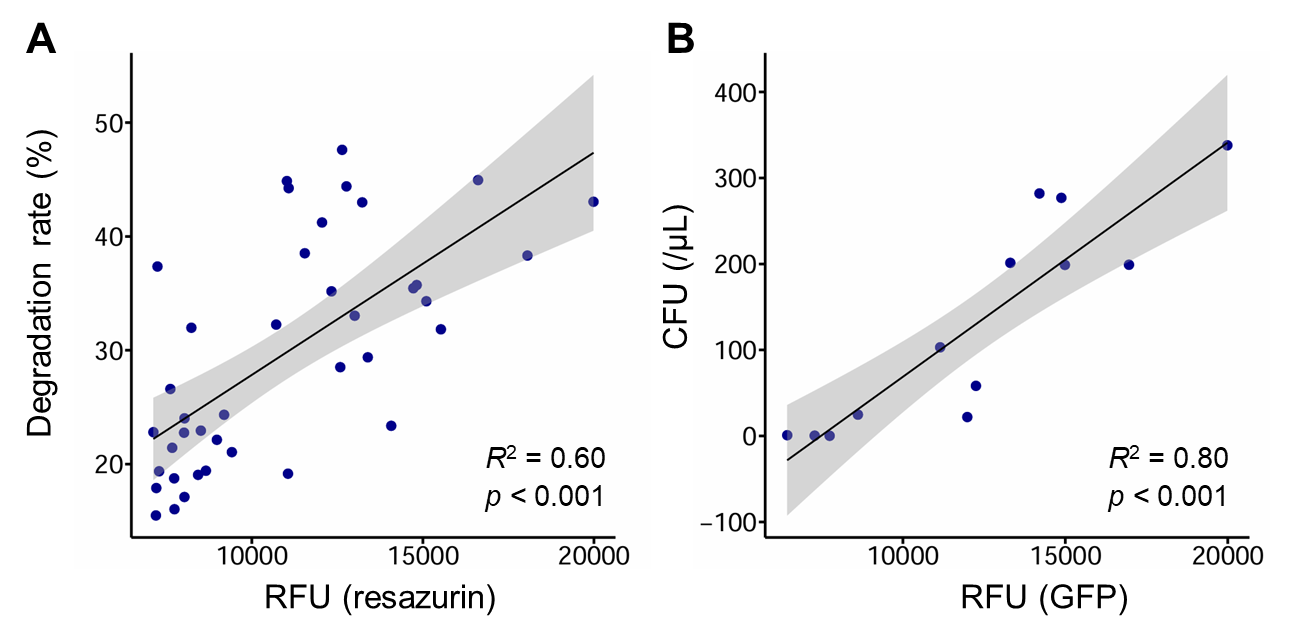
**Fig. S3** Validation of fluorescence-based measurements of aniline-utilization activity and Cab074F growth. Selected microplate cultures were evaluated for resazurin RFU and aniline degradation rate, or GFP fluorescence and colony forming units of Cab074F; paired values were compared (see Materials and Methods). A) Correlation between resazurin fluorescence and aniline degradation rate. B) Correlation between GFP fluorescence and CFU of strain Cab074F.

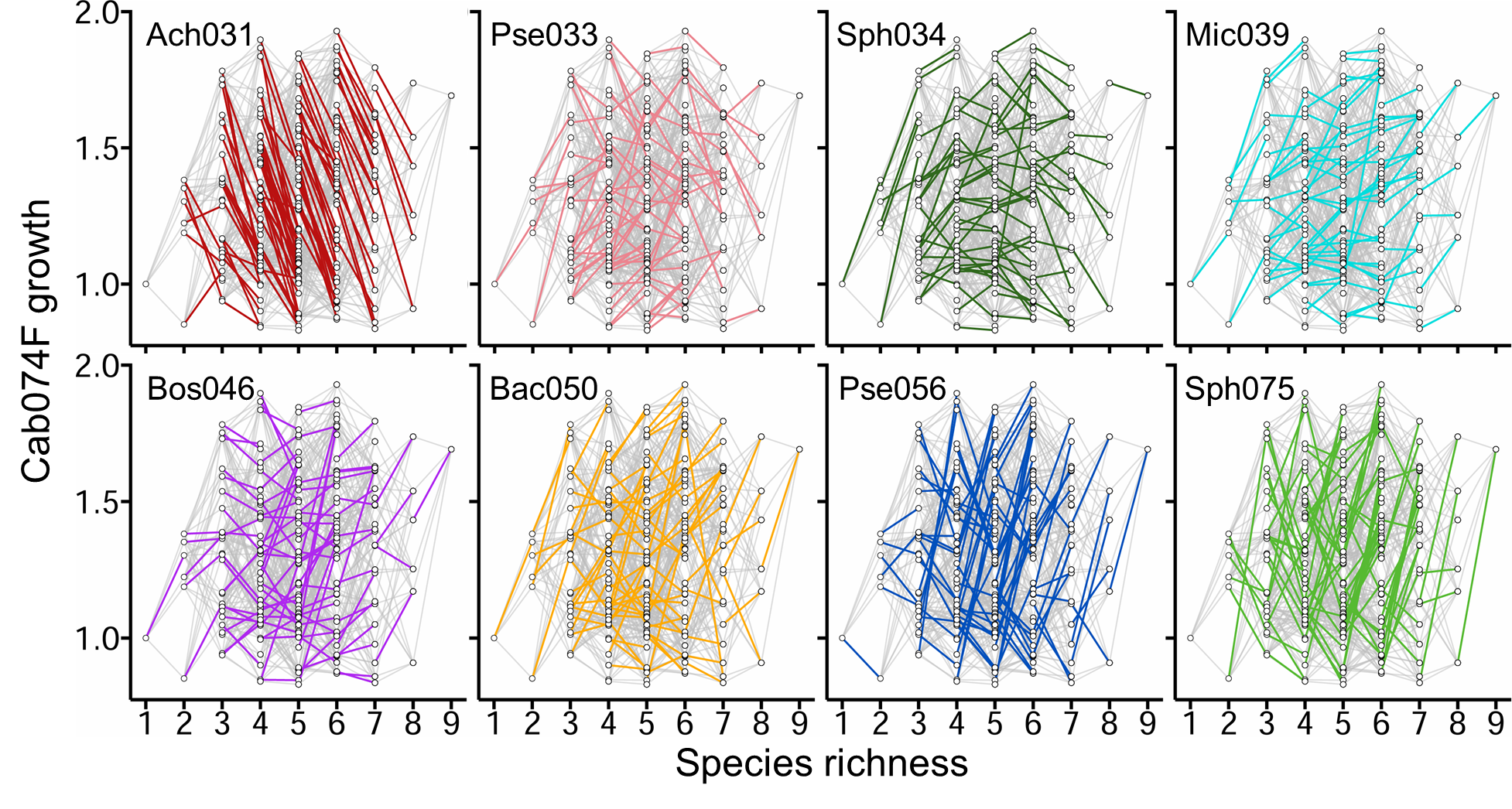
**Fig. S4** Composition–abundance landscape constructed from the dataset of strain Cab074F’s growth in the 256 sub-community combinations. Each data point represents a sub-community combination with a given species richness. The angles of the edges indicate the effect of each non-degrader strain on Cab074F growth in different community contexts.

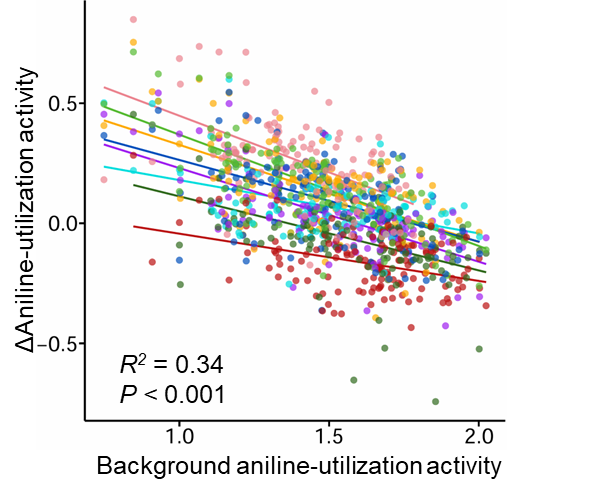
**Fig. S5** Correlation between the functional effect of each non-degrader strain and the functional level of the background community. The change in aniline-utilization activity upon adding a non-degrader strain (Δaniline-utilization activity) was plotted against the aniline-utilization activity of the corresponding background subcommunity combinations. Colors as in Fig. S4.

| Parameter | Operation conditions |
| --- | --- |
| Column | ODS-80Ts (Tosoh, Tokyo, Japan) |
| Mobile phase | CH_3_CN:H_2_O:CH_3_COOH = 400:600:1 |
| Detection wavelength | 240 nm and 260 nm |
| Flow rate | 0.5 mL/min |
| Injection volume | 20 µL |
| Column temperature | 40℃ |

**Table S1** Analytical conditions for HPLC

**Table S2** Identification results of synthetic community members based on 16S rDNA sequences. The closest MAG–isolate correspondence was predicted based on species-level identification results.

| Strain | Most similar species | Top-hit strain | Similarity | Closest MAG | Accession |
| --- | --- | --- | --- | --- | --- |
| Ach031 | *Achromobacter veterisilvae* | LMG30378 | 99.5% | MAG02 | LC878797 |
| Pse033 | *Pseudomonas monteilii* | NBRC 103158 | 99.9% | MAG03 | LC878798 |
| Sph034 | *Sphingomonas pseudosanguinis* | G1-2 | 99.4% | - | LC878799 |
| Mic039 | *Microbacterium laevaniformans* | DSM 20140 | 99.7% | MAG06 | LC878800 |
| Bos046 | *Bosea spartocytisi* | SSUT16 | 99.9% | - | LC878801 |
| Bac050 | *Bacillus mexicanus* | FSQ1 | 99.8% | - | LC878802 |
| Pse056 | *Pseudomonas guguanensis* | JCM 18416 | 97.8% | MAG05 | LC878803 |
| Cab074 | *Caballeronia cordobensis* | LMG 27620 | 99.7% | MAG01 | LC878804 |
| Sph075 | *Sphingomonas cannabina* | DM2-R-LB4 | 98.3% | MAG04 | LC878805 |

**Table S3** Quality, abundance, and taxonomic placement of metagenome-assembled genomes (MAGs). Completeness and contamination were assessed with CheckM. Abundance refers to the proportion of metagenomic reads mapping to each MAG. The closest species and average nucleotide identity (ANI), computed with DFAST, are also shown.

| ID | Completeness (%) | Contamination (%) | Abundance (%) | Closest species | ANI (%) |
| --- | --- | --- | --- | --- | --- |
| MAG01 | 94.8 | 0.4 | 37.8 | *Caballeronia cordobensis* | 97.7 |
| MAG02 | 98.1 | 0.4 | 27.9 | *Achromobacter denitrificans* | 99.2 |
| MAG03 | 80.7 | 0.0 | 9.9 | *Pseudomonas monteilii* | 96.6 |
| MAG04 | 99.7 | 0.9 | 1.2 | *Sphingomonas kyeonggiensis* | 89.3 |
| MAG05 | 97.0 | 0.6 | 0.7 | *Pseudomonas insulae* | 86.9 |
| MAG06 | 99.5 | 1.0 | 0.5 | *Microbacterium laevaniformans* | 94.0 |
| MAG07 | 94.4 | 0.7 | 0.5 | *Pseudorhodoferax aquiterrae* | 88.8 |
| MAG08 | 91.8 | 0.6 | 0.4 | *Stenotrophomonas nitritireducens* | 92.3 |
| MAG09 | 98.3 | 1.9 | 0.3 | *Sphingobium fuliginis* | 89.4 |
| MAG10 | 89.0 | 0.0 | 0.2 | *Methylophilus methylotrophus* | 83.3 |
| MAG11 | 87.3 | 0.0 | 0.2 | *Methylophilus rhizosphaerae* | 80.4 |
| MAG12 | 51.7 | 0.0 | 0.1 | *Acinetobacter pittii* | 97.1 |
| MAG13 | 84.0 | 0.7 | 0.1 | *Microbacterium hydrocarbonoxydans* | 82.9 |
